# COSTS AND BENEFITS OF ACTING TOGETHER: A NON-HUMAN PRIMATE MODEL OF COOPERATIVE DECISION-MAKING

**DOI:** 10.1101/2025.11.20.689048

**Authors:** Eros Quarta, Irene Lacal, Stefano Grasso, Andrea Schito, Virginia Papagni, Alexandra Battaglia-Mayer

## Abstract

Cooperation is a crucial aspect of social behavior, allowing individuals to achieve goals unattainable alone. However, collective actions require costly inter-individual coordination, making continuous cost-benefit evaluation essential before engaging in cooperation. Non-human primate models can help uncover the evolutionary foundations of human cooperation and its underlying neural mechanisms. To this aim, we developed a novel paradigm to analyze the dyadic behavior of two macaques, who took turns in choosing between individual and cooperative actions to obtain variable rewards. Each monkey used a joystick to guide a cursor toward one of two targets, each indicating both the reward magnitude and the action type (i.e. ‘solo action’ or ‘joint action’) required to obtain their payoff. We first observed a linear improvement in dyadic performance with the expected reward magnitude. Although macaques tended to prefer individual actions, they selected to act jointly under favorable payoff conditions, indicating that voluntary cooperation in macaques can emerge as a reward-driven process. Logistic models of their choices revealed the subjective cost monkeys assigned to cooperation, which was consistent across subjects. Nonetheless, the gain rate across different sessions increased over time, suggesting that macaques possess the cognitive ability to optimize their dyadic strategies, by estimating not only the coordination costs but also the benefits of cooperation. We observed a progressive reduction in the subjective cost assigned to joint action, which was not directly dependent on performance improvements. Our findings suggest that non-human primates can weigh the costs and benefits of cooperation, highlighting their ability to dynamically adjust social strategies based on reward contingencies.

**HIGHLIGHTS:**

- Expected reward magnitude enhances cooperative performance in macaques
- Monkeys evaluate the cost-benefit trade-off of cooperative actions in social contexts
- In macaques, the subjective cost of cooperation decreases over time, promoting a gradual shift toward a pro-social strategy that maximizes outcomes
- Cooperative behavior in macaques can emerge as a reward-driven, cost–benefit decision process
- Macaques may exhibit metacognitive dynamics, using internal evaluations of expected outcomes to guide collective behavior

## INTRODUCTION

Cooperation is fundamental for animal survival and thriving as it enables individuals to achieve goals unattainable alone. In joint actions (JA), a form of cooperation, agents must integrate internal models of self and others’ actions in real time (Stoit et al., 2011; Liebermann-Jordanidis et al., 2021). Such complexity is faced through coordination strategies that agents enact to achieve their shared goals, including reduction of individual variability and inter-individual differences in spatio-temporal aspects of motor behavior, which have been observed both in humans (Vesper et al., 2011; Satta et al., 2017) and non-human primates (Visco-Comandini et al., 2015; Lacal et al., 2022). During cooperation, humans might benefit from a ‘we-mode’ mental setting (Gallotti & Frith, 2013; Sacheli et al., 2018; Kourtis et al., 2019), allowing agents to be aware of their relative contributions, with an a priori sense of doing something together, rather than independently from each other. Interestingly, a similar ‘we-representation’ appears to emerge in macaques as well, when they engage in coordinated dyadic actions (Lacal et al., 2022). To act in synergy with another agent is considered costly due to the modest control subjects have on the partner’s intentions, actions and the resulting outcome (Wolpert et al., 2003; Gordon et al., 2021; van der Wel et al., 2021; Russo et al., 2025; Sacheli, Grasso et al., under review). On the other hand, acting with others can at times make a task easier, like transporting an heavy object (Wahn et al., 2018) and enabling goals otherwise precluded to a single agent, such as producing a complex symphony (Keller et al., 2014). Moreover, a costly cooperation can become worthwhile in high-stakes situations. For instance, a student would be more willing to work in a team, despite the cost of coordinating their actions with others, if a jointly performed exam offered more credits than if it was performed individually. In this framework, we consider decisions and ensuing actions during goal-directed social exchange as emerging from the integration of costs and benefits of a given option before selecting an appropriate action type, akin to deliberating between goods under various degree of uncertainty (Padoa-Schioppa, 2011; De Petrillo & Rosati, 2021). Crucially, while choosing between actions, human agents consider at least some aspects of the actions per se (Cos et al., 2011; Carsten et al., 2023) and such embodied decision-making seems to guide choice behavior also in macaques (Jun et al., 2021).

With specific regard to cooperation, it is therefore essential to evaluate the cost of coordination in relation to potential benefits that may arise, when deciding whether to act alone or jointly with others in a social context. From this perspective, macaques constitute an ideal model to study the neural basis of such processes, as they are capable of adapting their behavior to the demands imposed by the partner for successful joint performance (Visco-Comandini et al., 2015; Ferrari-Toniolo et al., 2019a; Lacal et al., 2022). Thus, as in humans, joint action in macaques entails coordination costs, as indicated by the changes in their kinematics parameters, such as shortened reaction times, elongation of movement times, reduced speed and straighter movement trajectories (Visco-Comandini et al., 2015; Satta et al., 2017; Lacal et al., 2022; Sacheli, Grasso et al., under review). These costs are also reflected in decreased probability of success when acting together compared to acting alone, likely arising from the uncertainties related to the partner’s intentions and higher complexity of motor control required to coordinate actions with a partner. At the neural level, this animal model revealed a correlate of joint motor interactions, through the identification of a class of premotor neurons, i.e., ‘joint-action cells’, preferentially active when monkeys act jointly with a partner (Ferrari-Toniolo et al., 2019a). In similar experimental settings, modulation of the Local Field Potentials (LFPs) was found to predict the effectiveness of interindividual coordination (Sacheli, Grasso et al., under review). In parallel, studies have shown that during social interactions monkeys integrate the partner’s behavior when making economic decisions (Haroush & Williams, 2015; Unakafov et al., 2020; Moeller et al., 2023).

Here we developed a behavioral paradigm that combines the decision-making aspects of cooperative behavior with those related to motor coordination required for joint actions, with the aim of laying the groundwork for an animal model to study the underlying neural mechanisms at single cell and population level. Here, we test the extent to which macaques are capable of performing cost-benefit evaluation when deciding between action types, namely acting alone or together. To address this, we analyzed the action type biases in their decisions alongside movement kinematics, allowing us to infer the subjective value monkeys assigned to cooperative actions.

## RESULTS

### Cooperative performance increases linearly with payoff

Two male macaque monkeys (Mk1, ≈ 9 kg; Mk2, ≈ 10 kg weight) were trained in a value-based decision-making task, to test their abilities in cost-benefit evaluation of cooperation when acting through joint actions (see Methods). The animals, seated side-by-side, choose between two offers and execute cursor movements, through an isometric joystick, towards the chosen target, each associated with an action-type and a pre-cued amount of liquid reward (water droplets). The task consisted of trials presented in random order across different choice contexts. The main condition of interest was the ‘action-type choice’ (ATC) trial type, in which monkeys, alternating the role of ‘decider’, were required to choose between solo action (SA) or joint action (JA) to receive a reward. Specifically, in each ATC trial, the ‘decider’ chose between two targets, each cued with a required action type (SA or JA) and a specific reward magnitude. Thus, the animals alternated the role of ‘decider’ (when called to make the above choice) and ‘responder’ (with potential engagement in JA) in a random order. These trials were defined as SA_i_–JA trials, with index *i* = 1, 2 indicating which monkey acted as the ‘decider’, choosing between its solo or joint action (**Fig. 1A–D**, Methods, and Movies 1, 2).

**Figure 1.**
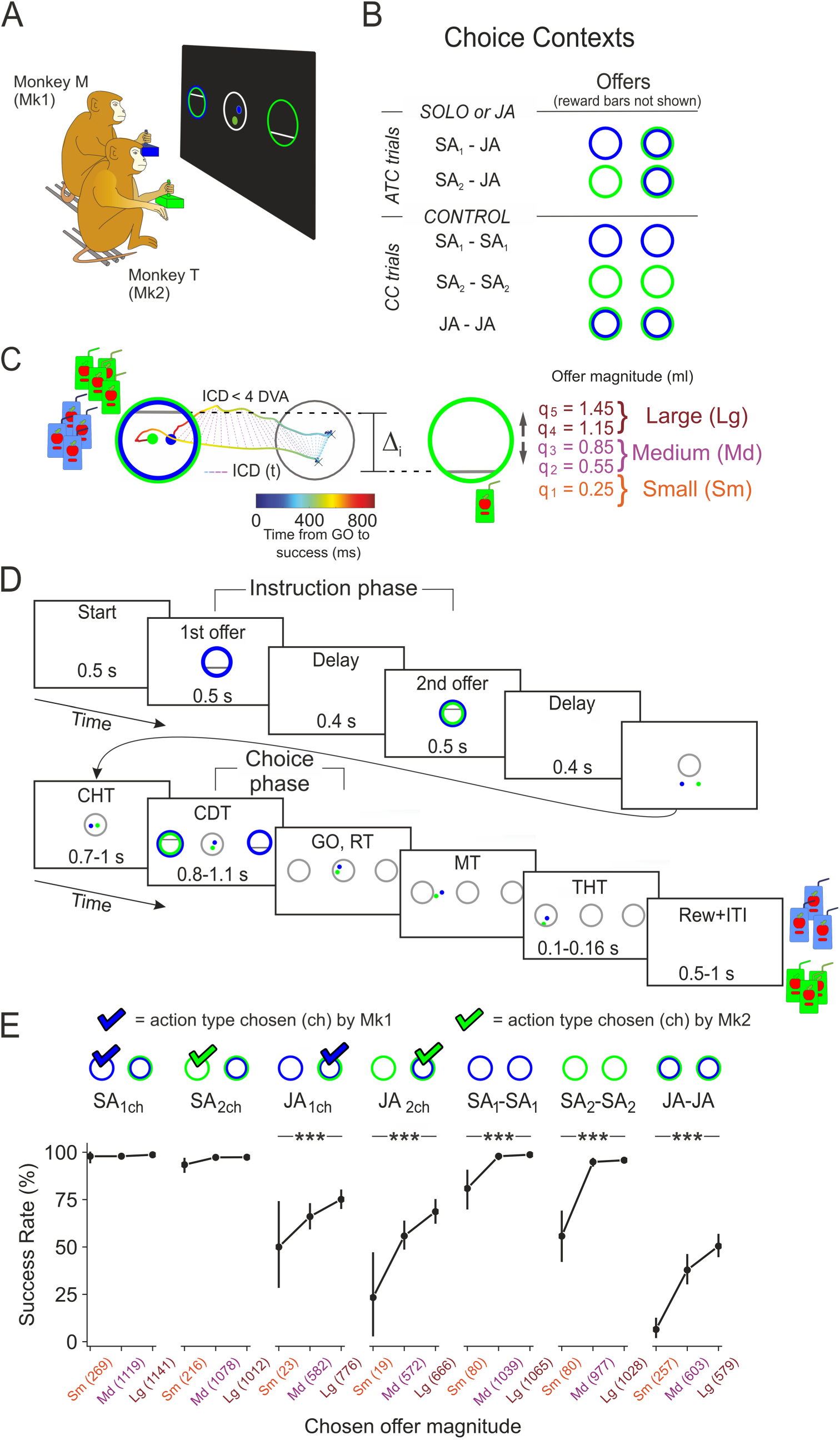
Experimental set-up and effect of expected reward on success rates. **(A)** Two monkeys sat side by side in front of a common monitor. Each monkey guided a visual cursor (blue, Mk 1; green, Mk2) from a central target toward a peripheral target by exerting a force on an isometric joystick to obtain a liquid reward. In each trial, two offers were presented, associated to (i) different action types, i.e. solo or joint action and (ii) varying reward magnitudes. **(B)** The monkeys were tested in two ‘action-type choice’ contexts (ATC trials) in which either Mk1 or Mk2 acted as the ‘decider’, choosing between solo or joint action (SA1-JA, SA2-JA trial), and three ‘control choice’ contexts (CC trials), in which the monkeys could choose between two offers obtainable through a given action type (SA1-SA1, SA2-SA2, and JA-JA trials). **(C)** Successful JA required the animals to coordinate their forces to bring jointly the two cursors into the peripheral target (bicolored green-blue circle), by keeping an inter-cursor distance (ICD) below 4 DVA to obtain both the reward. Cursors’ trajectories and ICD as a function of time (t) are shown for a typical trial in which monkeys choose the JA offer. Black crosses indicate cursors’ positions at movement onset. Horizontal gray bars cue the monkeys about the payoff size associated with each offer. Absolute offer magnitude (qi, i=1,2,3,4,5) ranged from 0.25 to 1.45 ml, resulting in 5 levels of reward differences Δi (i=0,1,2,3,4). The offer amounts were classified as Small (Sm: q1), Medium (M: q2, q3), and Large (L: q4, q5). **(D)** In each trial the two offers were first displayed sequentially in a central position, and after each monkey placed the cursor inside a central target (center holding time; CHT), they were presented simultaneously on the left and on the right side at 8 DVA from the center. After a choice delay time (CDT) the offers disappeared, and two gray circles served as a GO signal to move the cursor(s) toward the chosen offer (Movement Time, MT). Once inside the target, the cursor(s) had to be kept there for a brief target holding time (THT) to obtain the chosen reward (Rew). Inter-trial interval (ITI) lasted 500-1000 ms. **(E)** The reward magnitude had a linear effect on the cooperation outcome, as measured by the success rates (Binomial proportion test: *** p<0.001). Only significant differences in performance between Sm and Lg trials are shown. The number of trials for the different offer amount is reported in parentheses.

The task also included Control Choice (CC) trials in which the monkeys chose between two offers differing only in reward magnitude, while the action context remained constant. Specifically, decisions and subsequent actions were performed under identical action contexts (SA_1_, SA_2_, or JA), resulting in three types of CC trials: SA_1_-SA_1_, SA_2_-SA_2_ (also indicated as SA_i_-SA_i_, i=1, 2) and JA-JA. In these choice conditions, either Mk1, Mk2, or both monkeys together, were responsible for selecting and reaching the target to obtain the associated reward (**Fig. 1A–D**, Methods, and Movies 3, 4).

We first examined how the chosen offer, i.e., the expected reward for a successful action, affected the performance across different choice contexts. In both humans and non-human animals, reward magnitude is known to influence behavior during individual (solipsistic) actions, enhancing performance (Mogami & Tanaka, 2006; Pegg et al., 2021) and paradoxically impairing it under high-stakes conditions (Smoulder et al., 2021). However, the role of expected reward in cooperative performance in macaques has not been explored. To address this point, we partitioned trials within each choice context in 3 groups based on the chosen reward magnitude (Small, Sm; Medium, Md; Large, Lg; see methods), and found a context-dependent effect of reward amount on performance (**Fig. 1E**). In SA_i_-JA trials, two patterns emerged: when monkeys chose to act alone, reward had no impact, likely due to a ceiling effect, as for SA success rates (SR) already exceeded 93% in Sm trials (**Fig. 1E**); at variance, when choosing JA, performance increased from Sm to Lg rewards in both subjects (on average from 50% to 75% in Mk1 and from 23% to 68% in Mk2; binomial proportion test, Benjamini-Hochberg corrected, p<10^-10^; **Fig. 1E**). In CC trials, expected reward scaled SR in both SA and JA contexts: individual performance increased from about 80% to 99% in Mk1 and from 55% to 96% in Mk2; in JA–JA trials, performance improved from 6% to 50% (binomial proportion test, Benjamini-Hochberg corrected, p<10^-10^; **Fig. 1E**). These findings demonstrate that expected reward magnitude has a strong influence on the outcome of cooperative behavior in macaques, with high reward significantly enhancing the probability of a successful performance.

### Coordination cost of joint action is reflected in reduced movement vigor

Looking at the differences in performance across action contexts in CC trials, where no decision about action type was required, we found that the SR of both monkeys was near optimal during actions performed individually in both animals, with Mk1 displaying slightly better performances than Mk2 (SA_1_-SA_1_: 97%; SA_2_-SA_2_: 93%; Kruskal–Wallis, Dunn’s post-hoc test with Benjamini-Hochberg correction, p<0.001 **Fig. 2A**). In contrast, SR dropped significantly in JA-JA trials to about 37% (p < 10^-10^; **Fig. 2A**). This marked decrease, consistent with our previous findings in similar joint action tasks (Visco-Comandini et al., 2015; Ferrari-Toniolo et al., 2019a; Lacal et al., 2022), likely reflects a ‘coordination cost’ that monkeys may take into account, particularly when they have the option to act alone or together. Furthermore, in these CC trials, both monkeys exhibited movements with slower speed profiles during joint compared to solo actions (**Fig. 2B**), as indicated by reduced peak speed (SA_i_-SA_i_ vs JA-JA, Mk1: p < 10^-10^; Mk2: p < 10^-10^; Mann-Whitney-Wilcoxon test two-sided; **Fig. S1A, Table S1**). This resulted in a suppressive effect on movement vigor during joint action, as measured by the inverse of the sum of RT and MT (Mann-Whitney-Wilcoxon test two-sided, SA_i_-SA_i_ vs JA-JA, Mk1: p < 10^-10^; Mk2: p < 10^-10^, **Fig. 2C**; **Table S1**). The vigor reduction was due to longer movement duration (Mk1: 418±75 ms in SA_1_-SA_1_ vs. 586±119 ms in JA-JA trials, p < 10^-10^; Mk2: 486±102 ms in SA_2_-SA_2_ vs. 587±111 ms in JA-JA, p < 10^-10^, Mann-Whitney-Wilcoxon test two-sided; **Fig. S1A**). Both monkeys reduced their RT significantly when acting together (Mk1: 233±65 ms in SA_1_-SA_1_ vs. 215±95 ms in JA-JA; Mk2: 272±120 ms in SA_2_-SA_2_ vs. 234±112 ms in JA-JA trials, Mk1: p < 10^-10^; Mk2: p < 10^-7^ Mann-Whitney-Wilcoxon test two-sided; **Fig. S1A**). The changes in RT and MT observed in the present value-based decision-making task are consistent with previous findings in monkeys performing joint actions (Visco-Comandini et al., 2015; Ferrari-Toniolo et al., 2019a; Lacal et al., 2022). They support an inter-individual coordination model in which shorter, less variable RTs facilitate action synchronization, and therefore coordination, with a partner. In contrast, elongation of MTs observed during joint behavior likely reflects increased action control demands of inter-individual coordination. Together, these factors increase the action cost when achieving goals through joint performance.

**Figure 2.**
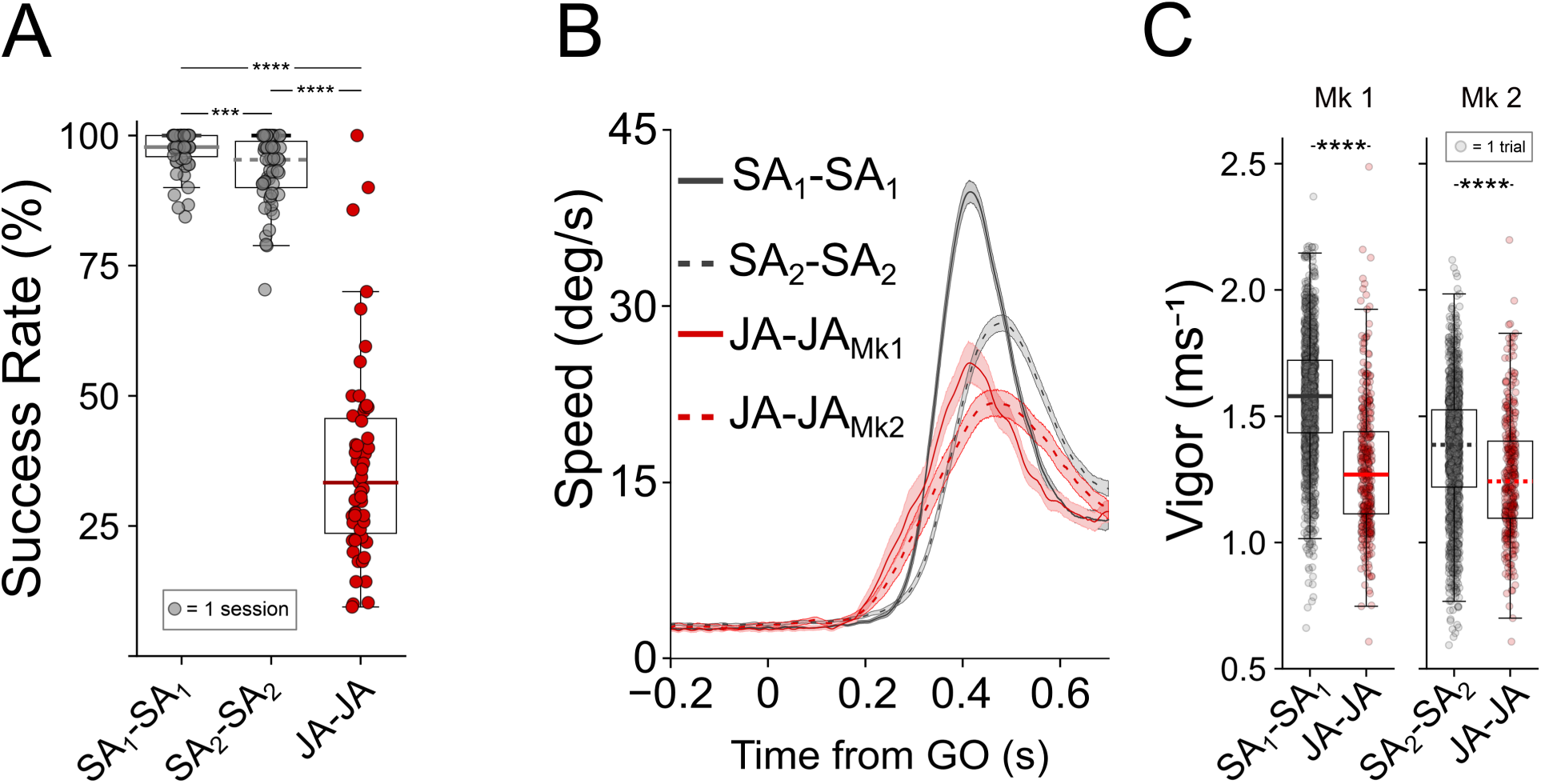
Performance during CC trials. **(A)** Percentage of success rates across choice contexts in CC trials: cooperating was associated to a significant decrease in performance, compared to individual actions (SA1-SA1, n=1579 trials; SA2-SA2, n=1471 trials; JA-JA n=702 trials). **(B)** Speed profiles aligned to the GO signal for each monkey during SAi-SAi (i=1,2) trials and JA-JA trials. Note the decrease in the peak speed for both monkeys when they acted together. **(C)** Movement vigor decreased during cooperation. Mann-Whitney-Wilcoxon test: ***: p <= 0.001, ****: p <= 0.0001.

### Context-dependency of cooperative performance

When analyzing the above parameters (SR, force application speed and Vigor) during ATC trials, i.e., when monkeys chose between different action types (SA or JA), we found as expected behavioral changes in line with those reported for CC trials (**Fig. 3**, **Table S2)**. Specifically, with respect to individual actions, following joint action choices i) performance dropped significantly (Kruskal–Wallis, Dunn’s post-hoc test with Benjamini-Hochberg correction, Mk1; p<10^-10^; Mk2, p<10^-7^, **Fig. 3A**); ii) force application speed was reduced (**Fig. 3B)**, as indicated by reduced peak speed (SA_i ch_ vs JA_i ch_, Mk1: p < 10^-10^; Mk2: p < 10^-10^; Mann-Whitney-Wilcoxon test two-sided; **Fig. S1B, Table S2**); and iii) movement vigor was also significantly reduced (Mann-Whitney-Wilcoxon test two-sided, Mk1: p=8.42×10^-243^; Mk2: p=1.38×10^-53^, **Fig. 3C**).

**Figure 3.**
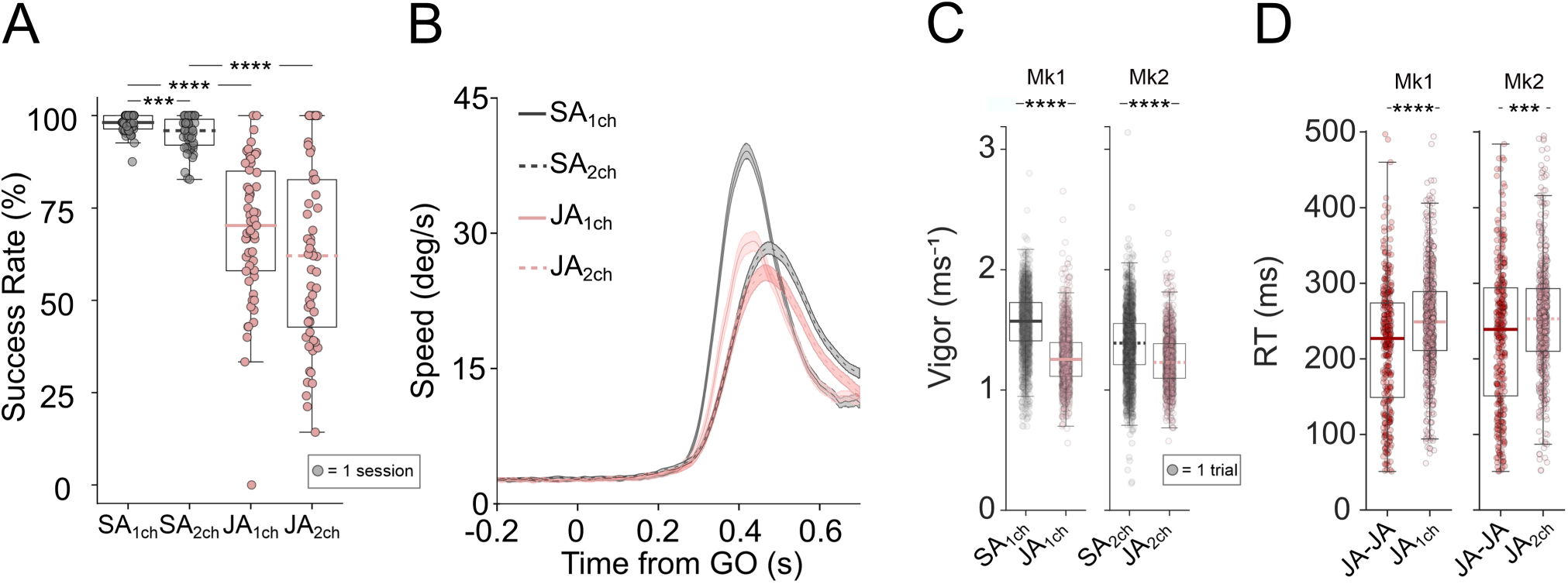
Context-dependency of cooperative performance. **(A)** Percentage of success rates across choice contexts in ATC trials (SAi-JA): choosing to cooperate is associated with a reduction in success rates compared to when monkeys chose to act alone. The index ‘i ch’ (i=1,2) specify the chosen action type by Mk1 or Mk2 during SAi-JA trials (SA1ch n=1878 trials; SA2ch n=1707 trials; JA1ch n=1519 trials; JA2ch n=1120 trials). (**B**) Speed profiles aligned to the Go signal for each monkey during ‘SA chosen’ and ‘JA chosen’ trials. Note the decrease in the peak speed for both monkeys when they chose to cooperate. ( **C**) Movement vigor decreased when monkeys chose to cooperate. (**D**) Comparisons of RTs associated with JA across choice contexts, i.e. during JA executed without explicit deliberation (JA-JA) or during JAi,ch (deliberated JA in SAi-JA trials). Mann-Whitney-Wilcoxon test: ***: p <= 0.001, ****: p <= 0.0001.

Moreover, we also observed behavioral changes when monkeys acted jointly after a deliberate choice to cooperate (SA_i_–JA trials), compared to JA–JA trials, in which such a choice was not required (**Fig. 3D, Fig. S1C**). In particular, the RT increase in trials requiring a choice between different action types (SA vs. JA) likely reflects the additional cognitive demands associated with evaluating and selecting between alternative action strategies prior to movement initiation, when choosing between the left/right target (**Fig. 3D**). The elongation of RTs was in line with the observed significant improvement in SR during joint action after active deliberation to cooperate (69% in ‘chosen JA_1_’ and to 62% in ‘chosen JA_2_’), compared with JA–JA trials (36% in JA–JA trials, Kruskal–Wallis, Dunn’s post-hoc tests with Benjamini–Hochberg correction, both p < 0.001; **Fig. 2A, 3A**). No changes in goodness of performance were observed when deciding to act alone relative to CC (SA_i_-SA_i_) trials. Taken together, these results suggest that the active deliberation of cooperation enhances joint performance, even though with no overall effect on movement vigor (**Fig. S1C**).

### Choice accuracy across action contexts

The behavioral modulation observed above across action types and/or choice contexts provides clear evidence that macaques incur a coordination cost when engaging in cooperative actions. Moreover, the reward sensitivity of both individual and joint action performance suggests that the decision to cooperate or act alone may be guided by the relative value of available offers.

While the behavioral and neural mechanisms of value-based decisions in the context of actions performed individually have been under intense scrutiny across economics and life sciences (Samuelson, 1947; Padoa-Schioppa & Assad, 2006; Glimcher & Fehr, 2013; X. Chen & Stuphorn, 2015; Reppert et al., 2015; Yamada et al., 2018; Zimmermann et al., 2018; Ballesta et al., 2020), such mechanisms have so far remained largely unexplored in the context of cooperating macaques. To fill this gap, we first characterized subjective choices strategies during trials that required monkeys to decide between two offers associated with the same action type, i.e., either acting individually (SA_i_-SA_i_ trials) or jointly with the partner (JA-JA trials). Mean choice accuracy values in SA_i_-SA_i_ trials were comparable to previous observations (Shi et al., 2022) on economic decision-making in non-human primates acting alone (Mk1: η=8.24±3.47; Mk2: η=7.14±4.5), indicating that they discriminated well between economic offers (see **Fig. S2A** for an example session). In JA-JA trials, accuracy (Mk1: η=8.03±3.47; Mk2: η=9.14±4.5) was comparable to that of SA_i_-SA_i_ trials, in both animals (Mann-Whitney-Wilcoxon test, SA_1_-SA_1_ vs JA-JA: p=0.3; SA_2_-SA_2_ vs JA-JA: p=0.5, two-sided with Bonferroni correction, **Fig. S2B**). Concomitantly, acting together was associated with increased variability of choice accuracy (Levene’s test, Mk1: p=1.5×10^-3^; Mk2: p=0.015, **Fig. S2B**). These latter results indicate that on average monkeys discriminated equally well between offers when acting alone or jointly and that choice accuracy was more variable when cooperating, possibly reflecting the uncertainty of joint decisions in the JA-JA trials.

Other variables being equal, monkeys engaged in individual actions might show a choice preference towards offers appearing on either side of the screen, defined as ‘side bias’ (Ferrari-Toniolo et al., 2019b). In our experiment, monkeys exhibited divergent side bias during SA execution: Mk1 had a left side bias, while Mk2 displayed an opposite preference towards offers on right side of the screen (Mk1: ρ_Side_=0.9±0.2; Mk2: ρ_Side_=1.5±0.5. Mann-Whitney-Wilcoxon test, U=176.0, p < 10^-10^, two-sided with Bonferroni correction. **Fig. S2C**). This result should not be overlooked, since it implies that to act together efficiently, monkeys had to find strategies to account for this divergency, either by reducing both their own bias or with one monkey adapting to the partner. We found evidence for the latter, as the side bias when acting together in Mk1 shifted towards the right side (JA-JA_Mk1_ trials: ρ_Side_=1.4±0.7; SA_1_-SA_1_ vs JA-JA_Mk1_ trials, Mann-Whitney-Wilcoxon test, U=800.0, p=2.1×10^-5^, two-sided with Bonferroni correction, **Fig. S2C**). No significant change was observed for Mk2 (JA-JA_Mk2_ trials: ρ_Side_=2.0±1.2; SA_2_-SA_2_ vs JA-JA_Mk1_ trials, Mann-Whitney-Wilcoxon test: U=1207.0, p=0.1, two-sided with Bonferroni correction, **Fig. S2D**). Thus, during JA, subjects had to consider individual idiosyncrasies, such as individual biases, that increased outcome uncertainty and reduced the probability of obtaining the reward (**Fig. 2A, 3A**).

### Evaluating the subjective value of cooperation in macaques

Once established that monkeys in CC trials discriminate equally well between offers and display a side bias when choosing between them, either when acting individually or together, we characterized their choice policy in ATC trials, i.e. when choosing to act alone or together. In a preliminary phase of our investigation (*prePhase*), in SA_i_-JA trials both subjects consistently preferred the SA, regardless of the relative difference in reward magnitude between the two offers. Even at the highest reward disparity (Δ4, corresponding to 0.60 ml), the choice rate for the JA offer remained below 25% (**Fig. S3**). We then hypothesized that although the reward for cooperation was highest in relative terms, its absolute quantity may have been insufficient to compensate for the coordination cost associated with joint action.

Thus, we tested the same monkeys in a new version of the task, where we doubled the unit of reward difference between the two offers (from *u* = 0.15 ml to *u* = 0.30 ml, see Methods). This manipulation produced a visually identical Δ4, which now corresponded to 1.2 ml, rather than 0.60 ml. In this new reward scenario, the action type choice pattern in ATC trials abruptly changed for both monkeys (**Fig. 4A**), with monkeys more prone to cooperate. Notably, the above reward settings were adopted for experimental data acquisition (*expPhase* sessions), and all reported analyses refer to these sessions. Logistic modeling revealed that the indifference point ρ_Coop_ in both animals was around Δ1 (∼ 0.3 ml, Mk1: ρ_Coop_=2.11±0.74; Mk2: ρ_Coop_=2.13±0.84; **Fig. 4B**). We considered the ρ_coop_ value as a proxy for the subjective cost that monkeys assigned to cooperation.

**Figure 4.**
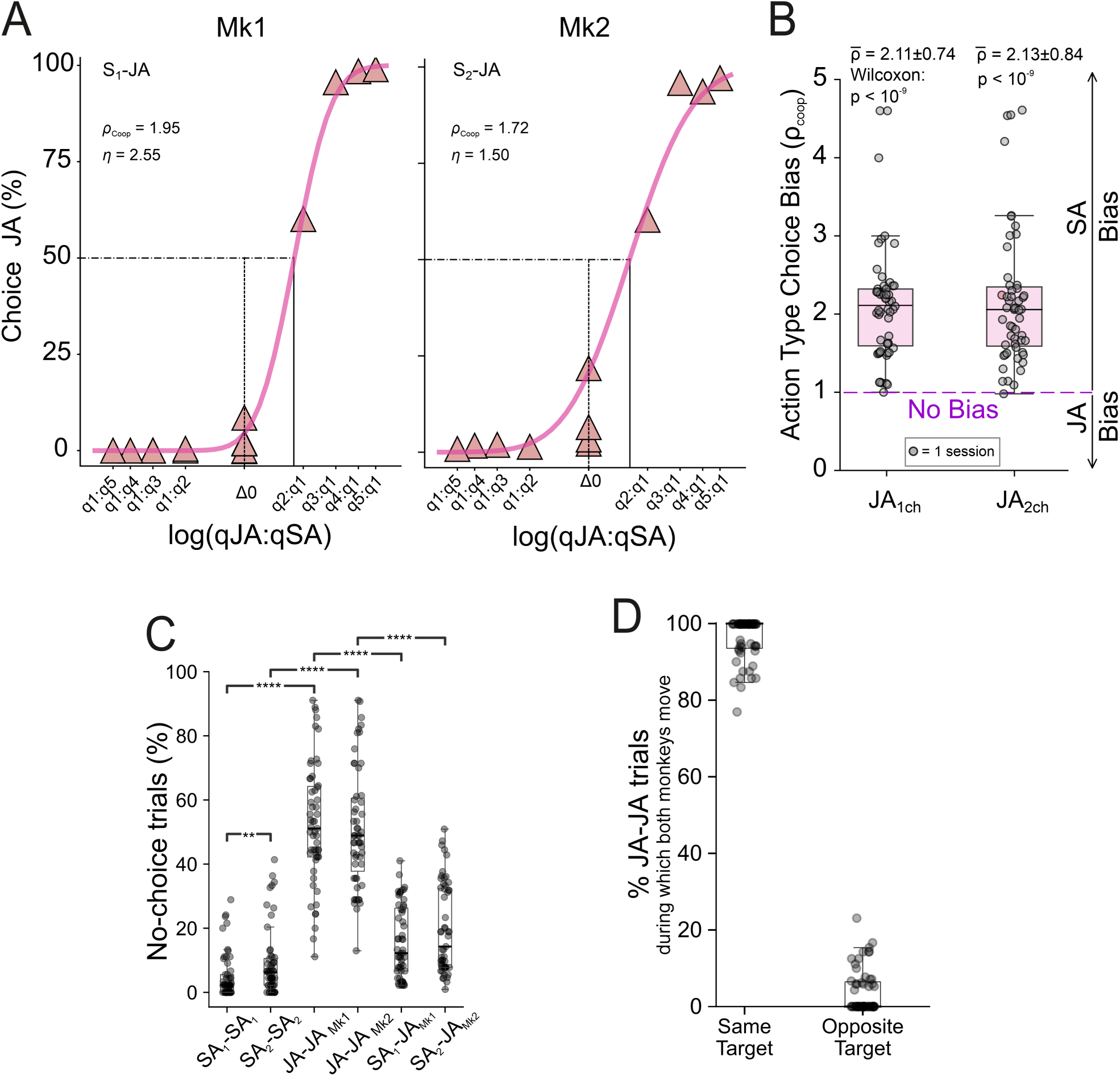
Cost-benefit evaluation when monkeys decide to cooperate or not. **(A)** Percentage of choices in favor of cooperation against the log quantity ratio between the two offers for JA vs SA_i_. Results refer to data pooling across experimental sessions (*expPhase*). Each data point represents percentages of ‘JA chosen’ trials for a given offer pair. Sigmoids were obtained from probit regressions. The relative value (ρ _Coop_) and sigmoid steepness (η) are reported. Note how the choices are biased in favor of the ‘solo’ action type ( ρ _Coop_ > 1). **(B)** Distribution of the ρ_Coop_ values across all sessions confirming the stark preference of both monkeys for the individual action type (Wilcoxon’s and t-test tests). **(C)** Percentage of ‘no-choice’ trials, i.e. monkeys prefer to stay in the center (decision avoidance), rejecting the offer. This percentage was markedly higher during JA-JA trials compared to solo trials (SA_i_-SA_i_), or when the decider monkey was required to choose between two different action types (SA_i_-SA_i_). Mann-Whitney-Wilcoxon test two-sided test: ***: p <= 0.001, ****: p <= 0.0001. **(D)** In JA-JA trials, when the animals decided to make a choice, they tended to always select the same target, thus making a congruent joint decision.

If choices between SA and JA actions were based solely on reward value, with no cost difference between the two, the indifference point would occur at Δ0 (i.e., ρ_coop_ = 1). Values of ρ_Coop_ > 1 instead indicate a preference for individual (SA) actions. Across sessions, ρ_Coop_ values were significantly larger than 1 (Mk1: Wilcoxon test p = 1.11×10^-11^; Mk2: Wilcoxon test p = 1.72×10^-10^; **Fig. 4B**), indicating the different action cost assigned by both animals to JA. Remarkably, in JA-JA decision context, monkeys most often adopted a ‘decision avoidance’ strategy, deliberately remaining in the center, effectively losing the possibility to obtain a reward (Mann-Whitney-Wilcoxon test two-sided, SA_i_-SA_i_ vs JA-JA, Mk1: p < 10^-10^; Mk2: p < 10^-10^, **Fig. 4C**). However, in JA-JA choice context, when restricting the analysis to trials in which both animals committed to a target, the two partners consistently selected the same option, resulting in a congruent joint decision (**Fig. 4D**). Interestingly, when the two targets were associated with different action demands (SA_i_–JA trials), and therefore with different efforts, the decision process appeared to be facilitated, as shown by the higher percentage of choices made under this choice condition (Mann-Whitney-Wilcoxon test two-sided, JA-JA vs SA_i_-JA, Mk1: p < 10^-10^; Mk2: p < 10^-10^, **Fig. 4C)**. These findings highlight the lower subjective value that macaques assign to cooperation, as reflected in their frequent disengagement during mandatory JA–JA trials and their systematic bias toward individual actions in SA_i_–JA trials. Importantly, the latter was neither entirely dictated by the offer nor by their action type preference, but rather based on a trade-off between the absolute amount of offered reward and subjective value each monkey assigned to cooperation.

### Temporal dynamics of subjective cost-benefit trade-off of joint action

Since in sensory decision-making choice strategies can vary continuously even in well-trained animals (Roy et al., 2021), we tested the temporal dynamics of choice rates in favor of joint actions in the ATC contexts, when monkeys were asked to decide between acting alone or together (SA_i_-JA). In both subjects, the designated ‘decider’ showed an increasing tendency to cooperate across sessions (Mann–Kendall test: Mk1, τ = 0.49, p = 7.9×10^-6^; Mk2, τ = 0.42, p *=* 0.006; **Fig. 5A, Fig. S4B**). This trend was accompanied by a robust correlation between the partners’ choice rate for JA (Kendall’s τ = 0.62 at lag = 0, p = 3.43×10^-11^, **Fig. 5A**), suggesting that reciprocal influences shaped their cooperative choices within each session. This growing joint propensity to cooperate raised the question of whether or not it was driven by improved joint performance over time. Interestingly, the monkeys did not show any progressive improvement in their joint performance (indexed by SR) across sessions when they decided to act together. If anything, performance exhibited a decreasing trend across sessions, which was statistically significant for Mk2 (Mann–Kendall test, Mk1: τ = −0.14, p = 0.122, Mk2: τ = −0.29, p = 0.006, **Fig. 5B**). We reasoned that, across sessions, monkeys began to attribute a lower cost to joint actions and increasingly engaged in higher-risk decisions, accepting greater probability to fail but with greater potential benefit. To test this aspect, we examined the temporal evolution of the subjective cost (ρ_Coop_) of joint action and choice accuracy (η) over sessions. In both animals, we observed a significant decrease in ρ_Coop_ (Mann–Kendall test; Mk1: τ = –0.64, *p* = 3.04 × 10^-6^; Mk2: τ = –0.61, *p* = 2.24 × 10^-4^; **Fig. 5C, E**) suggesting a progressive lower cost assigned by both animals to JA over time. Similarly, we found a significant decrease in η (Mk1: τ = –0.43, *p* = 6.01 × 10^-4^; Mk2: τ = –0.61, *p* = 0.01; **Fig. 5D, E**), indicating greater variability in the animals’ choices and less deterministic decision boundaries, likely reflecting an increased tendency toward exploratory behavior favoring joint action (JA). Although both ρ_Coop_ and η showed visually similar temporal trends between animals, Kendall’s τ indicated weak positive correlations (τ = 0.18 for ρ_Coop_, τ = −0.19 for η), neither of which reached statistical significance (p = 0.057 and p = 0.056, respectively; Fig. 5C, D).

**Figure 5.**
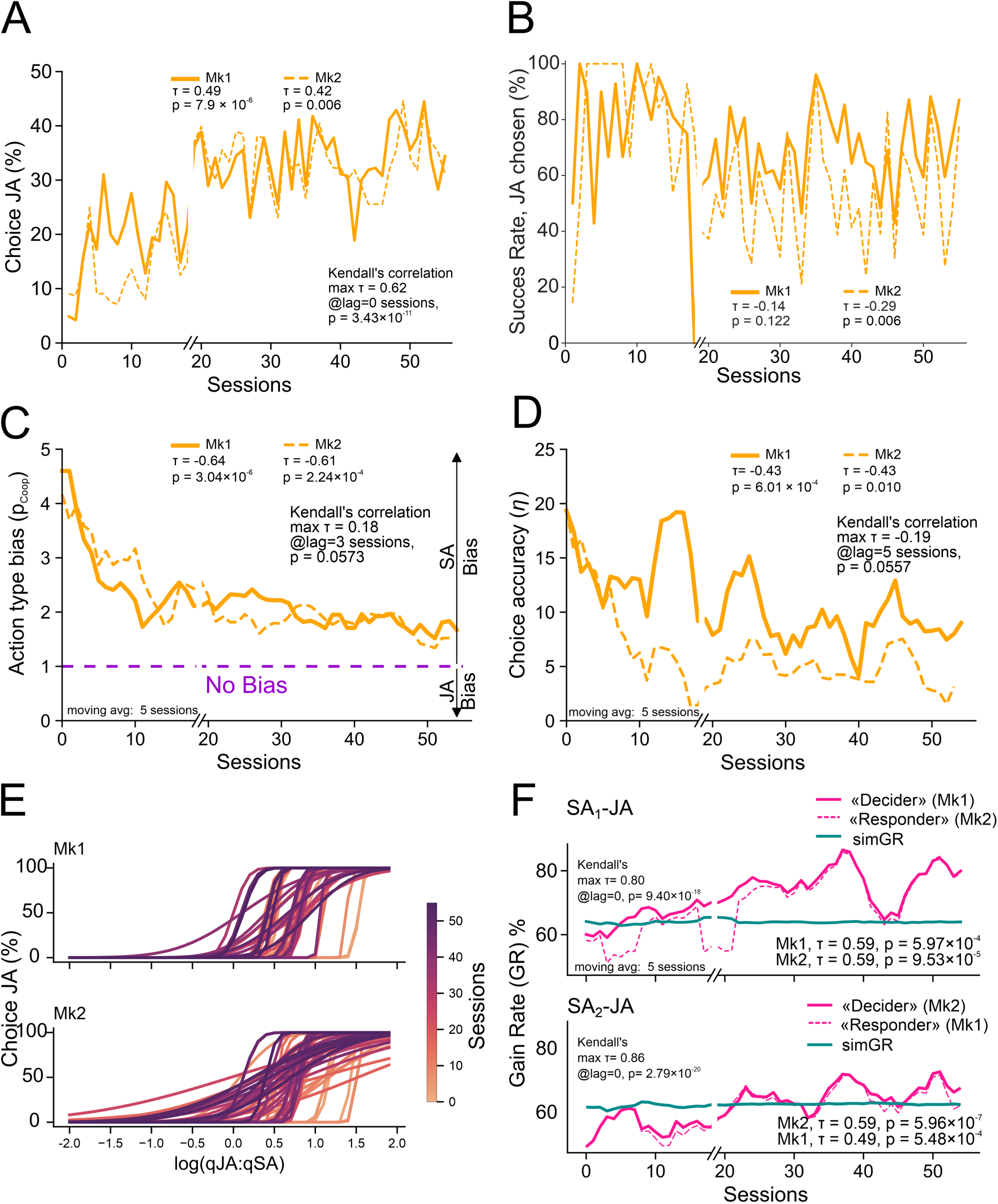
Monkeys gradually enforce a pro-social choice policy that increases their global gain rates. **(A)** Choice rate in favor of cooperation increased across sessions in both monkeys. (**B)** Time-resolved success rate associated with JA in SAi-JA choice context (’JA chosen’ trials). **(C)** Both monkeys displayed a decrease in ‘action-type bias’ across sessions (Kendall’s τ<0, p<.001). **(D)** Choice accuracy decreased across sessions, suggesting increasing variability in their choice strategies. **(E)** Psychometric curves of all individual sessions for SAi-JA trials are plotted for each monkey and color-coded based on their temporal order (session number). Note that the initial sessions (orange) are shifted toward the right side of the x-axis, indicating a stronger bias in favor of individual action, compared with the later sessions (violet), which also display a shallower slope. **(F)** In SAᵢ–JA trials, the gain rate (GR) was computed as the percentage of reward collected relative to the total amount obtainable in each session. GR increased significantly across sessions for both monkeys (Kendall’s τ > 0, p<.001; ‘decider’, solid magenta; ‘responder’, dashed magenta), indicating optimization of reward income through a progressively more pro-social choice policy. In each plot, the experimental GR was compared to the simulated GR (simGR, green solid line) corresponding to the percentage of reward per session that would have been obtained if the animals would have exclusively chosen to act alone (SA offer). The significant positive trends suggest that over time the monkeys learned the benefits of cooperation.

In light of the observed temporal trend in the propensity to cooperate, a key question concerns the underlying positive drivers of this increasing tendency toward joint action. To address this, we analysed reward outcomes in SA_i_-JA trials by measuring, for each session, the amount of reward actually gained by the animal (either as a ‘decider’ or ‘responder’) and compared it to the hypothetical reward they would have obtained, had they consistently chosen the SA offer. Specifically, we computed the Gain Rate (GR), defined as the ratio between the total reward collected by the animal in SA_i_-JA trials and the sum of the highest available offers across those trials (see Methods). We then compared this to the simulated GR (simGR), which estimates the reward that the animal would have received by systematically selecting the individual option. We were interested in the time-varying dynamics of the GR across sessions. Indeed, we found that when either monkey was the ‘decider’, both animals increased their own GR (Mann-Kendall test, SA_1_-JA: Mk1 τ = 0.59 p = 5.97×10^-4^, Mk2 τ = 0.59 p = 9.53×10^-5^; SA_2_-JA: Mk2 τ = 0.59 p = 5.96×10^-7^, Mk1 τ = 0.49 p = 5.48×10^-4^, **Fig. 5F**). Thus, the GR emerging from a pro-social choice policy integrating cooperation with solo actions, increased over time. These findings show that macaques are capable not only to accurately evaluate the costs of coordination, but also the benefits of collaborative actions, and over time select optimal dyadic strategies to maximize their reward.

## DISCUSSION

Social interactions rely on an internal cost-benefit computation: is the effort of coordination worth the potential gain from cooperation? Understanding this trade-off is critical for analysing the mechanisms that support social behavior. We focus here on contexts where individuals can achieve greater rewards via joint actions but must also invest effort in precise motor coordination with a partner.

In evolutionary biology and behavioral economics, a cooperator is defined as someone who pays a cost (*c*), so that another individual receives a benefit (*b*) (Nowak, 2006). In the context of motor interaction, coordinating one’s own action plan with that of a partner to accomplish a common objective, otherwise unattainable by acting alone, can be viewed as the cost *c* to be paid. Here and in the motor domain, ‘coordination’ is used referring to the spatio-temporal alignment necessary to achieve a shared goal (Sebanz & Knoblich, 2009).

To investigate the evaluation processes underlying cost-benefit trade-offs in social decision-making, we used a non-human primate (NHP) model, which enables us to study the evolutionary roots of human cooperative behavior, especially where action-type decision-making and associated motor planning are integrated. While cooperation (Haroush & Williams, 2015) and coordination (Ong et al., 2021; Moeller et al., 2023) in macaques have been examined in game-theoretical frameworks emphasizing cognitive and social aspects of such processes, our approach leverages on the motor aspects of dyadic interactions and focus on how macaques evaluate the costs of cooperation planned and executed through joint actions. Our experimental paradigm provides a platform for exploring the neural computations underlying the trade-off between coordination costs and cooperative gains. We devised a two-alternative choice paradigm in which pairs of macaques could choose between acting alone or coordinating with a partner to obtain varying reward magnitudes. This design allowed us to estimate the subjective coordination cost for each animal and track how this cost evolved over time. Previous work shows that macaques can flexibly adjust their motor plans and execution strategies to meet the demands of joint action (Visco-Comandini et al., 2015; Lacal et al., 2022). Here, we demonstrate that such flexibility extends to decision processes: macaques not only adapt their motor behavior when acting with a partner, but also adjust strategic choices over time to optimize reward, engaging in joint action, only when advantageous. Rational behavioral updating has recently been reported in chimpanzees (Schleihauf et al., 2025). Our findings suggest that macaques may exhibit similar metacognitive dynamics, using internal evaluations of expected outcomes to guide collective behavior, through mechanisms that ultimately enhance survival in dynamic social environments.

### Reward influence on cooperative performance

Expected rewards are known to drive behavioral performance (Schultz, 2000; Berridge, 2004; Kobayashi et al., 2006; Schultz, 2015; Watabe-Uchida et al., 2017; Smoulder et al., 2021) and guide the allocation of cognitive resources in effortful tasks (Frömer et al., 2021). In NHPs performing individual actions, moderate reward magnitudes promote more vigorous and accurate responses (Smoulder et al., 2021). We show that this principle extends to joint actions in macaques: dyadic performance improved linearly with reward magnitude. Moreover, higher rewards enhanced the animals’ willingness to engage in joint action.

Regarding the NHPs’ attitude to cooperate, when behavior is studied in macaques through game-theory approaches like the iterated Prisoner’s Dilemma (Haroush & Williams, 2015), reluctance to cooperate was observed. In the wild, whether NHP, particularly great apes, can intentionally coordinate to achieve otherwise inaccessible goals remains debated (Melis, 2013). Controlled laboratory studies have shown that great apes possess the coordination skills for collaboration and can recruit partners, when necessary (Melis et al., 2006), but unlike humans as young as three years old (Hamann et al., 2011) are often reluctant to share rewards (Melis & Tomasello, 2013). This lack of prosocial motivation is believed to prevent the emergence of more complex, human-like cooperative behavior, by diminishing the intrinsic incentive to engage in joint efforts. Also macaques, like great apes, when studied under highly controlled lab contexts show motor coordination skills for joint action (Visco-Comandini et al., 2015; Ferrari-Toniolo et al., 2019a; Lacal et al., 2022). Here, we add that they can also voluntarily engage in cooperation under adequate motivational conditions, increasing cooperative choices when joint rewards are sufficiently high. Crucially, the motivation to cooperate seems to be not merely driven by relative payoff differences between individual and joint commitments. While in trials where both offers required the same action type (SA_i_–SA_i_ or JA–JA), animals consistently chose the higher-reward offers, regardless of absolute value or difference, a higher expected reward for joint-, as compared to solo-action, is not by itself sufficient to consistently promote cooperative behavior. This was true also when the reward difference was maximal. In other words, the best available option in relative terms was not always enough to elicit joint engagement. Only sufficiently high absolute rewards for joint action overcame the cost of coordination and justify its effort. These findings suggest that, as in great apes, macaques can assess the value of cooperative engagement in a quantitative and accurate fashion.

### Effort cost evaluation of JA behavior

Motor and choice behaviors are not only reward-driven, but also shaped by effort-based decision-making. The concept of effort, though variably defined, is generally perceived as costly and aversive (Inzlicht et al., 2025). Yet, effort is also valued in goal-directed behavior, and is often essential for optimal goal pursuit. Effort can be in general associated with both physical and cognitive costs. In the present experiment, monkeys were specifically required to exert greater cognitive control during joint actions. This demand is evidenced by the modulation of their movement kinematics, which aimed at optimizing dyadic performance, as previously documented in our studies (Visco-Comandini et al., 2015; Lacal et al., 2022). In the current study, this modulation was also reflected in a significant decrease in movement vigor during joint action compared with individual action. Such increased cognitive demand during joint action execution likely reflects the need for enhanced online motor control and continuous monitoring of the partner’s action, allowing the animal to flexibly adapt its behavior to that of the other agent.

This effort is also reflected in the decision-making phase when monkeys had to coordinate their choices jointly to direct their cursor in one of the two targets, such as in JA–JA trials. Compared to individual choices in the solo condition (SA_i_–SA_i_ trials), joint decisions in JA-JA trials were marked by a greater reluctance to choose in both animals, suggesting that the cost of coordination also impacts the decision processes, resulting in a lower probability to optimize reward achievement. While decision avoidance was most frequent when both options required joint performance (JA–JA trials), the decision process appeared to be facilitated when the two targets involved different action demands (SA_i_–JA trials), and thus more discriminable effort levels. In these cases, it was easier for the animals to assess which of the two rewards was worth the additional effort required by cooperation.

This brings us to the estimation of the cost monkeys attribute to inter-individual coordination compared to acting alone. As already mentioned, they showed a significant bias in favor of individual actions over joint actions, when they were free to choose how to obtain their reward. In SA–JA trials, the ρ_Coop_ value (action type bias) was used as a proxy for the subjective cost assigned to joint action. Interestingly, ρ_Coop_ values were highly similar in the two animals (**Fig. 4B**), suggesting that both assigned comparable coordination costs and shared similar expectations about the payoff required to justify the effort of their cooperation.

Our results align with the economic principle that decision-making seeks to maximize the difference between ‘utility’ (reward benefit) and ‘disutility’ (effort cost). Studies in humans (Croxson et al., 2009; Westbrook & Braver, 2015; Inzlicht et al., 2025) and other animals (Bautista et al., 2001) show that effort is treated as a cost weighed against potential rewards (Burrell et al., 2023). The principle holds across species, with choice behavior well explained by subtractive models of effort. In our task, effort was not manipulated experimentally but emerged from the two action types: individual actions involved lower effort than joint actions. While consistent with a subtractive discounting model, our two-level effort design precludes comparing alternative computational models (e.g., nonlinear functions).

At the neural level, inter-individual coordination in joint action is likely supported by a distributed fronto-parietal network, in which circuits encoding flexible eye–hand coordination provide a cortical substrate where motor control and decision processes can interact (Battaglia-Mayer, 2019). A recent study (Sacheli, Grasso et al., under review) revealed in humans stronger bilateral dorsal premotor and parietal BOLD activation during joint actions, with premotor activity correlating with the goodness of inter-individual coordination. The same study showed that, in macaques’ PMd, LFPs are not only modulated by joint action but also encode the level of interpersonal coordination.

### From costs to benefits: Temporal shifts in cooperative choice

Choice policies also evolved over time. Although sensory decision-making is often modelled with fixed psychometric functions, strategies can change even in well-trained animals (Roy et al., 2021). In value-based contexts, recent work shows temporal context effects shaped by expectations of overall effort shaped by past experience (Burrell et al., 2023). In our task, the proportion of cooperative choices in SA_i_–JA trials increased over sessions. Interestingly, the increased motivation to act jointly was not driven by improved performance, and therefore by a reduced coordination effort, but rather by a greater willingness to accept coordination risks for higher rewards. This shift corresponded to a progressive increase in accumulated rewards, surpassing what they would have earned by always acting alone.

In conclusion, our findings suggest that macaques can quantitatively evaluate the cost–benefit balance in social interactions, by weighing expected payoff against coordination effort. Like in great apes, they assess the value of cooperation and choose to collaborate when rewards outweigh costs. By integrating motor demands, cognitive control, and expectation of future rewards, macaques provide a valuable model for studying the neural and behavioral bases of simultaneous interplay of motor coordination and cooperative decision-making.

### Limitations of the study

Our findings emerge from a relatively restricted sample size (n=2 monkeys) and should be interpreted cautiously when generalizing to the cognitive processes underlying cooperation. At the same time, the dyadic performance and movement kinematics during control trials (SA_i_-SA_i_ and JA-JA trials) were in line with previous investigations of our laboratory, aggregating to n=6 monkeys, which is well beyond the standard sample sizes in neurophysiological experiments in macaques. Importantly, while some behavioral metrics were subject-specific, our new study revealed how the choice policies were remarkably similar between the two monkeys, hinting at fundamental rules of engagement for cooperation in the primate brain.

## ACKNOWLEDGMENTS

This study was supported by the Ministry of University and Research, Italy (grant nr. B53D23014490006 & B53D23030110001 to ABM, Young Researcher 2024 grant nr. B83C24003140006 to EQ) and by the Sapienza University of Rome (to ABM and EQ). We thank Roberto Caminiti for his useful comments and suggestions during the preparation of the manuscript. We thank Denise Tomassetti for help with animal husbandry.

## AUTHOR CONTRIBUTIONS

ABM designed the experiment with contributions from IL. IL, EQ, SG, AS, and ABM collected the data. IL, EQ, and ABM designed the analysis. EQ and IL analyzed the data with contributions from SG, VP, ABM. EQ, IL, and ABM wrote the first draft of the article. All authors revised and approved the final article.

## METHODS

### ANIMALS AND EXPERIMENTAL SETUP

Two male *Macaca mulatta* monkeys participated in the experiment (Monkey M, ≈ 9 kg; Monkey T, ≈ 10 kg weight). Hereafter, the two animals are referred to as Mk1 and Mk2. All animal care and housing procedures complied with European (EU Directive 63/2010) and Italian (DL 26/2014) regulations on the use of non-human primates in scientific research.

The monkeys were placed in a soundproof chamber, seated side by side on two primate chairs facing a 40-inch monitor (100 Hz refresh rate; 800×600 resolution; 32-bit color depth; viewing distance: 150 cm; **Fig. 1A**). They were positioned at least 60 cm apart to prevent physical contact, and the orientation and structure of the chairs minimized direct visual contact.

Both monkeys were trained to control a visual cursor (diameter 0.6 degrees of visual angle, DVA) displayed on a black screen by applying dynamic force with the hand on an isometric joystick (ATI Industrial Automation, Apex NC, USA). The joystick sampled force in two dimensions at 1 kHz on a horizontal plane. The amount of force applied was proportionally converted into cursor displacement on the x–y axes of the vertical plane of the monitor, such that a force of 1 N moved the cursor from the central position to an eccentricity of 1.25° DVA. Both monkeys used their left arm to perform the task, based on their preference during early training, while the other arm was gently restrained. The cursor could be guided individually (solo action, SA) or jointly with the partner (joint action, JA), as described below. During joint action, the monkeys had to coordinate their forces in space and time to move their cursors into a shared target (bicolor, green–blue circle) without exceeding an inter-cursor distance (ICD) threshold of 4 DVA from the go signal to entering the chosen target (**Fig. 1C**). Successful coordination yielded a reward for both animals, whereas failure resulted in no reward.

Task control and behavioral data collection were managed with the NIH-funded software package REX. Eye movements of both animals were monitored using an infrared oculometer (Arrington Research), sampled at 1 kHz, and stored along with joystick force signals and key behavioral events.

### BEHAVIORAL TASK

The value-based decision-making visuomotor task used in the present study was designed to investigate choice behavior and movement kinematics in monkeys free to choose between two targets. Each target was associated with a specific reward magnitude and one of two possible action types required to obtain the reward: solo action (SA) or joint action (JA). The reward amount was indicated by a level bar inside the circular target, while its color specified the action type (SA₁ for Mk1: blue; SA₂ for Mk2: green; JA: bicolor green–blue; **Fig. 1**). The action types consisted of either moving a single cursor from the central position toward a peripheral target alone (solo) or coordinating cursor motion with that controlled by a partner (JA). Animal behavior was examined in a preliminary phase (*prePhase*) and an experimental phase (*expPhase*) using tasks with identical trial structures (see ‘Trial structure’) but differing in the reward magnitudes associated with the two offers (see ‘Reward magnitudes setting’). Each monkey controlled an isometric joystick to guide its cursor (Mk1: blue; Mk2: green) into the chosen target.

In each session, the two animals were tested in randomly interleaved decision-making trials belonging to five choice contexts (**Fig. 1B**), subdivided into action-type choice (ATC) and control choice (CC) contexts. In ATC trials, the monkeys alternated the role of ‘decider’, selecting one of two targets, each associated with a specific reward amount and an action type (SA or JA). These trials are referred to as SA_1_–JA and SA_2_–JA trials (or also as SA_i_-JA, i=1, 2 where the index indicates which monkey acted as the ‘decider’, choosing between a solo or joint action). While one animal decides between the two action types, its partner could either remain in the central target or act jointly, depending on the decider’s choice. The possible game dynamics were as follows: (i) the ‘decider’ chooses SA. The partner receives the same reward by remaining in the central target. Conversely, if the partner stays in the center, the ‘decider’ can only obtain a reward by acting alone toward the SA target. (ii) The ‘decider’ chooses JA. The only way for both animals to receive a reward is to cooperate by moving together to the same target.

In addition to ATC trials, we included CC trials (**Fig. 1B**) in which the animals chose between two offers differing only in reward magnitude while the action type was held constant. Three CC contexts were used: SA_1_–SA_1_, SA_2_–SA_2_, and JA–JA. In these contexts Mk1, Mk2, or both monkeys together, respectively, were responsible for selecting and reaching the target to obtain the reward. These control conditions allowed us to test whether the monkeys could discriminate between offers requiring the same action type. During SA_1_–SA_1_ (SA_2_–SA_2_) trials, Mk1 (Mk2) decided between the two targets while its partner could remain in the center to receive the same reward chosen by the ‘decider’. In the JA–JA choice context, the two monkeys were presented with two joint action offers and had to make a coordinated decision to move together into one of the two targets to obtain the indicated reward. No reward was delivered if they failed to reach the same target jointly and in a coordinated fashion.

#### Reward magnitude setting

As described above, in each trial the two offers were each associated with a specific reward magnitude. The reward consisted of a variable volume of water, indicated to the animals by a level bar displayed inside each target. We define the ‘absolute reward’ (*AbsRew*) as the reward magnitude associated with a given option, and the ‘relative reward’ (*RelRew*) as the difference between the magnitudes assigned to the two options. As noted above, the tasks adopted in the *prePhase* and *expPhase* differed only in their reward-setting criteria.

##### PrePhase task settings

To explore how behavior depended on reward, this preliminary version used a broader variability of reward assignments than the final task, within an arbitrary established range of 0.10 - 1.40 ml (see below). Specifically, the absolute amount of reward for each option was randomly defined on each trial under the following constraints: i) the absolute rewards fell within a range from 0.10 ml to 1.40 ml; ii) the relative amounts of reward (Δᵢ, with i = 0,1,2,3,4) between the two options were kept fixed at different values (Δ₀ = 0 ml; Δ₁ = 0.15 ml; Δ₂ = 0.30 ml; Δ₃ = 0.45 ml; Δ₄ = 0.60 ml. Accordingly, the payoff value for the two options was calculated as:

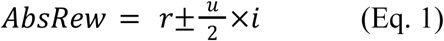

where r is a randomly assigned base reward amount, *u* is the unit of reward difference (*u* = 0.15 ml), and *i* is the delta coefficient (*i* = 0,1,2,3,4), implying

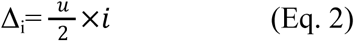

The task was programmed so that the higher reward was presented either as 1^st^ offer or as 2^nd^ offer (**Fig. 1D**), in a balanced way.

A preliminary analysis of the data collected with this ‘*prePhase*’ version of the task showed that the monkeys were: i) very weakly motivated to choose between the two offers, particularly during joint action conditions, frequently displaying decision avoidance and, as a result, often failing to complete the task; and ii) when asked to choose between acting alone or jointly, they almost never selected the joint action option, even under the most favorable relative-reward condition (Δ₄ = 0.60 ml), for which the proportion of trials in which the JA option was chosen fell below 25% (**Fig. S4**).

We therefore hypothesized that, although the relative reward for cooperation was highest, its absolute value may still have been insufficient to offset the coordination costs associated with acting together. To test this, we introduced a new version of the ATC trials in which we doubled the unit reward difference between the two offers (from *u* = 0.15 ml to *u* = 0.30 ml, see below). This modification led to a markedly different incentive scenario and was adopted as the final version of the experimental task. The number of trials collected with the *prePhase* task were 9314 across 28 sessions.

##### ExpPhase task settings

The final version of the task differed from the preliminary version in two aspects: i) the absolute reward was not randomized, but rather fixed; ii) the unit of reward difference *u* was doubled, from 0.15 ml to 0.30 ml. Therefore the amount of reward (*AbsRew*) associated to each offer was set as follows:

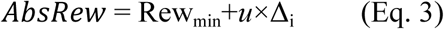

where *Rew_min_* is a fixed amount of the offered reward, *u* = 0.30 ml is the unit of reward difference in the final version of the task) and Δ_i_ (i=0,1,2,3,4) is the step size difference between offers (**Fig. 1C**), as defined above (Eq. 2).

The value of *Rew_min_* was calculated using behavioral data from the *prePhase* task settings. It was computed as the reward minimum amount needed to maximize the likelihood that monkeys would complete the trial, and thus make a choice between the presented offers. We estimated *Rew_min_* in the following steps: i) we collected the reward amounts from Δ₀ trials in which a choice was made; ii) we ordered those values in ascending order; iii) we computed the mean of the lowest 30% of values, separately for each ATC and CC condition. This method, rather than using the sharp minimum, was preferred since it is more robust against outliers. Finally, we selected *r_min_* as the higher value amongst the means computed for each choice condition.

In Δ_0_ trials (i.e. with relative difference between options equal to zero), the absolute reward levels were randomly selected from the following five possible amounts; (r*_min_*, r*_min_* + *u*×1, r*_min_* + *u*×2, r*_min_* + *u*×3, r*_min_* + *u*×4.

Because the unit size *u* was doubled compared with the *prePhase* task, the offer deltas in the present version of the task were also doubled, yielding the following values: Δ₀ = 0 ml; Δ₁ = 0.30 ml; Δ₂ = 0.60 ml; Δ₃ = 0.90 ml; Δ₄ = 1.20 ml, while absolute reward levels corresponded to: 0.25 ml (q1), 0.55 ml (q2), 0.85 ml (q3), 1.15 ml (q4), 1.45 ml (q5) and were cued through a level bar displayed on the two peripheral targets (**Fig. 1C**).

The number of trials collected in the final version (exp*Phase*) of the task, which were used for the full behavioral analysis, were 16902, obtained testing animal behavior in 55 sessions.

#### Trial structure

Each trial comprised two main phases: an ‘instruction phase’ and a ‘choice phase’ (**Fig. 1D**). Instruction phase: The two offers were first displayed sequentially (first and second offer) at a central, spatially neutral position on the screen. Each offer was presented for 500 ms, separated by a 400 ms delay. After both offers were presented, two cursors and a central gray target appeared. Each monkey was required to place its cursor inside the central target and hold it there for a center-holding time (CHT; 700–1000 ms).

Choice phase: the same two offers were then presented again, one on the left and the other on the right side of the central target (randomized across trials), 8 degrees of visual angle (DVA) from the center. After a choice-delay time (CDT; 800–1000 ms), the two offer targets, showing their color (action type) and level bar (reward magnitude), were turned off, leaving only two neutral gray circles. This event served as the GO signal for the monkeys to move their cursors from the central target toward their chosen peripheral target or to remain in the center. The ‘center stay’ was a rewarded option for the ‘responder’ in cases where the ‘decider’ acted individually toward the peripheral targets associated with its solo action. In such cases, the center stay provided the partner with the same reward amount obtained by the ‘decider’ acting in the solo condition.

The interval from cursor movement onset to the achievement of the final target, more precisely corresponding to the ‘dynamic-force time’ was, for simplicity, defined as ‘movement-time’ **Fig. 1D**), This interval had to be completed within 2 s, otherwise, the trial was aborted. After reaching the chosen target, the animal(s) had to maintain the cursor(s) on it for a short target-holding time (THT; 100–160 ms; **Fig. 1D**) to receive the chosen reward (Rew; **Fig. 1D**). Videos of reconstructed representative trials for each choice context are described in the supplementary information and provided as supplementary files (Movies 1-4).

### QUANTIFICATION AND STATISTICAL ANALYSIS

#### Success Rates

To analyze choice behavior of each monkey, a decision in a given trial was registered when the cursor exited the central circle following the ‘GO’ signal, towards one side. If the cursor did not fulfill the above condition the monkey’s behavior was classified as ‘no-choice’ for that specific trial. Using this procedure, the behavior of each monkey during every trial was classified as either ‘choice made’ or ‘no-choice’. The overall behavioral motor performance of the monkeys was quantified using the success rate (SR), defined for each choice context as the percentage of successful (i.e., completed) trials relative to the total number of trials in which the monkey made a decision in that context (i.e., we discarded from this analysis the ‘no-choice’ trials). Specifically for the JA-JA trials, we discarded those trials for which both monkeys stayed in the center. Statistical comparisons across contexts were performed using a Kruskal-Wallis Test followed by post-hoc pairwise comparisons with Dunn’s test and Bonferroni correction for multiple testing. To test the effect of reward on performance, for each choice context we grouped SR based on 3 levels of chosen offer: small (Sm, corresponding to quantity q1), medium (Md, corresponding to quantities q2 and q3) and large (Lg, corresponding to quantities q4 and q5). Given the relatively low number of trials, to compare performance across offer groups, we used a two-tailed binomial proportion test comparing Small-to-Medium, Small-to-Large, and Medium-to-Large conditions (Smoulder et al., 2021).

#### Movement kinematics

To investigate temporal features of movement planning when acting together, we analyzed the reaction times (RT), defined as the time elapsing from the go signal to the onset of the cursor’s movement; given that the cursor’s position was proportional to the force applied by the animal to the isometric joystick, it was the expression of the applied dynamic force. The onset of the cursor’s movement was defined as the time at which, for at least 90 ms, the cursor’s speed and the cursor acceleration exceeded by three SD the average speed and acceleration signal calculated in a 50 ms time-window before the presentation of the peripheral targets. Trials for which RT was shorter than 50 ms were also excluded from the analysis (Ferrari-Toniolo et al., 2019a). Movement Time (MT) was quantified as dynamic force time, defined as the time spanning from the onset of the cursor’s movement to its entry into the peripheral target. To obtain insights about the subjective value of acting together, we employed movement vigor, defined here as the inverse of the sum of RT and movement duration (Shadmehr et al., 2019; Bufacchi & Iannetti, 2021). The peak speed was calculated as the maximum value of the cursor’s tangential speed for each trial. To investigate the strength of dyadic coordination we measured the point-by-point distance between the two cursors controlled by the monkeys (inter-cursor distance, ICD) during the execution of the joint actions. To quantify the extent to which each monkey moved to minimize spatial deviations between starting and target entering positions, we computed the movement curvature as the ratio between the path length of the actual movement and the path length of a straight trajectory with the same start and ending positions.

#### Choice rates and modeling

To investigate the subjective value assigned by the monkeys to solipsistic and cooperative actions, choice patterns of the monkeys were modeled using logistic regressions. Across all trial types, the psychometric curve steepness measures the choice consistency and herein we adopt the term choice accuracy (Padoa-Schioppa, 2022). To quantify ‘side bias’, choice patterns during control trials (SA_i_-SA_i_, JA-JA) were modeled using probit regressions with the following equation:

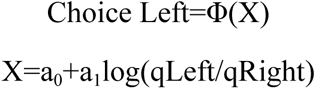

where choice Left=1 if the animal chose the offer on the left side and 0 otherwise, *Φ* is the cumulative function of the standard normal distribution, qLeft (qL) and qRight (qR) the quantities offered for the left side and for the right side offer, respectively (Padoa-Schioppa, 2022). To quantify choice accuracy and action type bias during SA_i_-JA trials, choices were modeled using probit regressions with a similar equation:

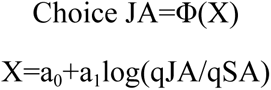

Here, Choice JA=1 if the animal chose the JA offer and 0 otherwise, *Φ* is the cumulative function of the standard normal distribution, qJA and qSA the quantities presented for the JA and the solo offer, respectively. Through data fitting we derived measures for the relative value

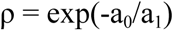

and the sigmoid steepness η = a1. The relative value ρ is the indifference point i.e., the payoff quantity that made the animal indifferent between the two options, therefore corresponding to the same probability to be chosen. In the case of SA_i_-SA_i_ and JA-JA trials the relative value ρ quantifies the *side bias*, while in SA_i_-JA trials ρ serves to quantify the *action type bias*, which is used as a proxy for the subjective cost of cooperation (ρ_Coop_). To test for potential long-term trends in choice accuracy/variability and subjective cost of cooperation, we applied a moving window average (window = 5 sessions) to the sequence of the η and ρ values across sessions to filter out higher-frequency fluctuations. From these time-series we extracted the Kendall’s tau (τ), and performed the Mann-Kendall non-parametric statistical test to quantify the strength and direction of monotonic trends (Chen et al., 2022).

#### Choice strategies

To analyze whether the monkeys’ choice strategies in SA_i_-JA trials were aimed at maximizing reward collection, we explored whether and to what extent the overall observable choices led to maximizing actual gain. To this end, we quantified the gain rate (GR) obtained during the experimental sessions by each monkey, defined as the ratio of the total quantity of reward collected by the subject in each session relative to the sum of each trial’s highest offer. We then compared the GR obtained from experimental data with a simulated gain rate that each animal could have gained if the ‘decider’ would have systematically chosen to act alone (simGR). To compute the latter, the SR were also simulated taking the levels achieved during the own solo or, in the case of the partner, the SR for staying in center while the ‘decider’ moved its cursor to the solo target. Mann-Kendall non-parametric statistical test was used to quantify the strength and direction of monotonic trends (Chen et al., 2022). The Mann-Kendall test was used to assess the presence of a monotonic trend across the ordered observations.

All analyses were conducted in MATLAB (MathWorks), Python, and R. Results are reported as mean±SD if not stated otherwise.

## SUPPLEMENTARY INFORMATION

### Movies

**Movie 1.** Video of a typical ‘action-type choice’ (ATC) trial (in this example SA_1_-JA) in which the ‘decider’ (Mk1) chooses to act alone (SA_1_ch) by guiding a blue cursor (session id MT022CH, trial 37), while the Mk2 acts as ‘responder’ staying in the center circle (green cursor) to obtain the same reward quantity. The red cross indicates the instant of movement onset (MoveON). The blue and green check marks signal the reward delivered to both monkeys. The red cross and the check marks are shown for illustrative purposes and not displayed to the monkeys. See **Fig. 1** and Methods for additional details.

**Movie 2.** Video of a typical SA_1_-JA in which the ‘decider’ (Mk1) chooses to act jointly (JA_1_ch) with the ‘responder’ (same session as in Movie 1, trial 239). See **Fig. 1** and Methods for details. Conventions and symbols as in Movie 1.

**Movie 3.** Video of a typical Control Choice (CC) trial (in this example SA_1_-SA_1_) in which Mk1 decides between two offers to be obtained by acting alone (same session as in Movie 1, trial 16). During this trial Mk2 has to keep its own cursor within the central target, until the end of the trial, to get the same reward quantity obtained by Mk1. See **Fig. 1** and Methods for details. Conventions and symbols as in Movie 1.

**Movie 4.** Video of a typical CC trial, in which monkeys were called to act together (JA-JA) in order to obtain their reward (same session as in Movie 1, trial 265). See **Fig. 1** and Methods for details. Conventions and symbols as in Movie 1.

**Table S1.** Movement kinematics during ‘control choice’ (CC) trials (mean±SD).

|  | Mk1 |  | Mk2 |  |
| --- | --- | --- | --- | --- |
|  | SA<sub>1</sub>-SA<sub>1</sub> | JA-JA | SA<sub>2</sub>-SA<sub>2</sub> | JA-JA |
| RT (ms) | 233±65 | 215±95 | 272±120 | 234±112 |
| MT (ms) | 418±75 | 586±119 | 486±102 | 587±111 |
| Peak Speed (dva/s) | 35.3±6.5 | 26.9±6.9 | 31.5±6.35 | 25.9±5.77 |
| Vigor (ms<sup>-1</sup>) | 1.57±0.23 | 1.30±0.26 | 1.37±0.24 | 1.26±0.24 |

**Table S2.**
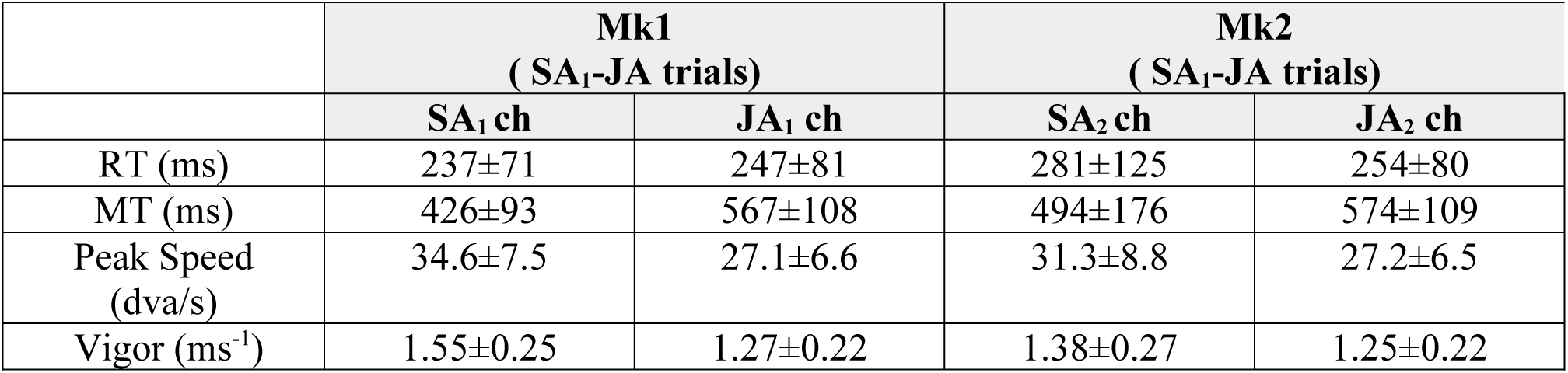
Movement kinematics during ‘action-type choice’ trials (mean±SD). The index ‘ch’ specifies the chosen action type by Mk1 or Mk2.

|  | Mk1<br>(SA<sub>1</sub>-JA trials) |  | Mk2<br>(SA<sub>1</sub>-JA trials) |  |
| --- | --- | --- | --- | --- |
|  | SA<sub>1</sub> ch | JA<sub>1</sub> ch | SA<sub>2</sub> ch | JA<sub>2</sub> ch |
| RT (ms) | 237±71 | 247±81 | 281±125 | 254±80 |
| MT (ms) | 426±93 | 567±108 | 494±176 | 574±109 |
| Peak Speed (dva/s) | 34.6±7.5 | 27.1±6.6 | 31.3±8.8 | 27.2±6.5 |
| Vigor (ms<sup>-1</sup>) | 1.55±0.25 | 1.27±0.22 | 1.38±0.27 | 1.25±0.22 |

**Figure S1.**
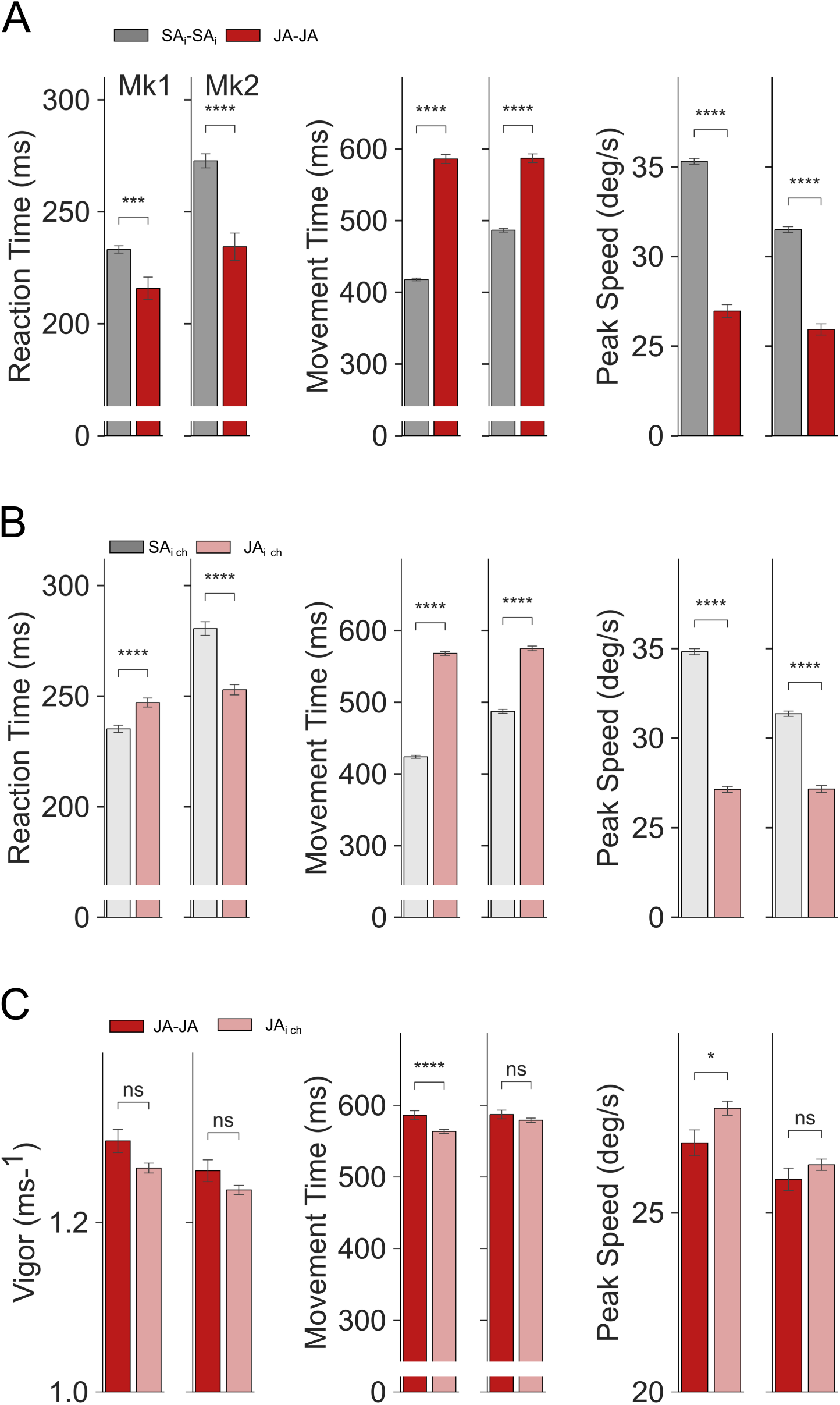
Movement kinematics across action types and choice contexts during successful trials. **(A)** Reaction times (RTs) were reduced when cooperating in JA-JA trials with respect to the SA_i_-SA_i_ trials. Concomitantly, movement times (MTs) were longer and the peak speeds were lower when cooperating vs when acting alone in these CC trials. **(B)** In ATC context, RTs increased when Mk1 chose JA compared to trials in which Mk1 chose SA. At variance, Mk2’s RTs when choosing to cooperate were lower compared to the trials in which this monkey chose SA. MTs were increased and peak speeds were reduced when choosing to cooperate with the other monkey in JA_i_ chosen (ch) trials. **(C)** Vigor, MTs and peak speeds when cooperating in CC and ATC trials. Monkeys displayed similar vigor when cooperating across contexts. MTs were reduced in Mk1 when choosing to act together, while this parameter was not changed in Mk2. Similarly, Mk1 increased its peak speed when choosing to act together, while this parameter was not changed in Mk2. Mann-Whitney-Wilcoxon test: *: p <= 0.05,***: p <= 0.001, ****: p <= 0.0001.

**Figure S2.**
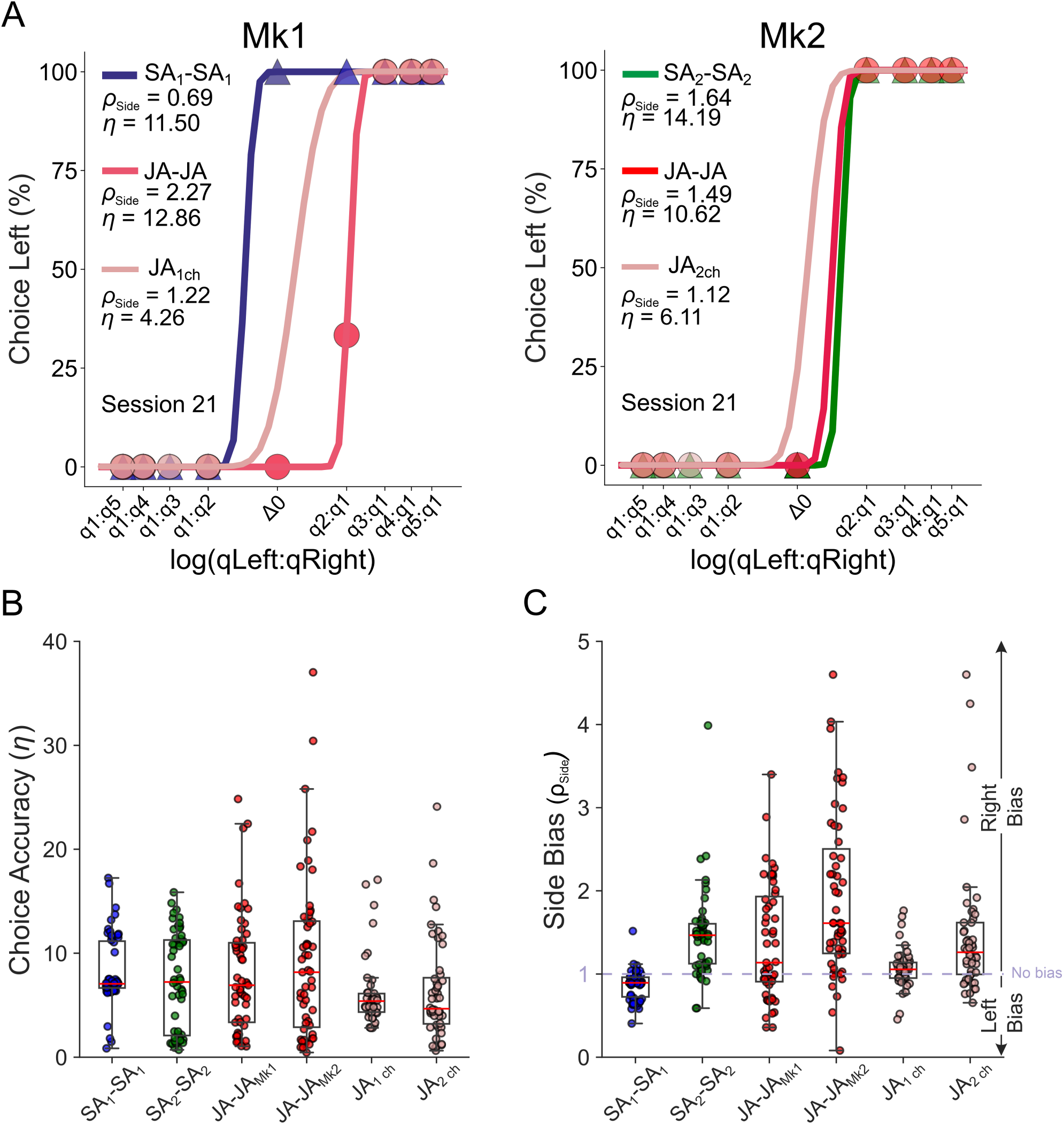
Side bias across action types and choice contexts. (**A)** The percentage of choices in favor of the left offer is plotted (y-axis) against the log quantity ratio between the two offers (left vs right, x-axis), for Mk1 and Mk2, for one typical session. Each data point represents percentages of “left chosen” trials for one offer pair. Sigmoids were obtained from probit regressions. The relative value (ρ_Side_) and sigmoid steepness (η) are reported as text. **(B)** Distribution of choice accuracy when acting alone or in cooperation with the other monkey. **(C)** Choices during SA_i_-SA_i_ trials are biased in favor of the offer presented on the left side for monkey 1 (ρ_Side_ < 1) and right side for monkey 2 (ρ_Side_ > 1).

**Figure S3.**
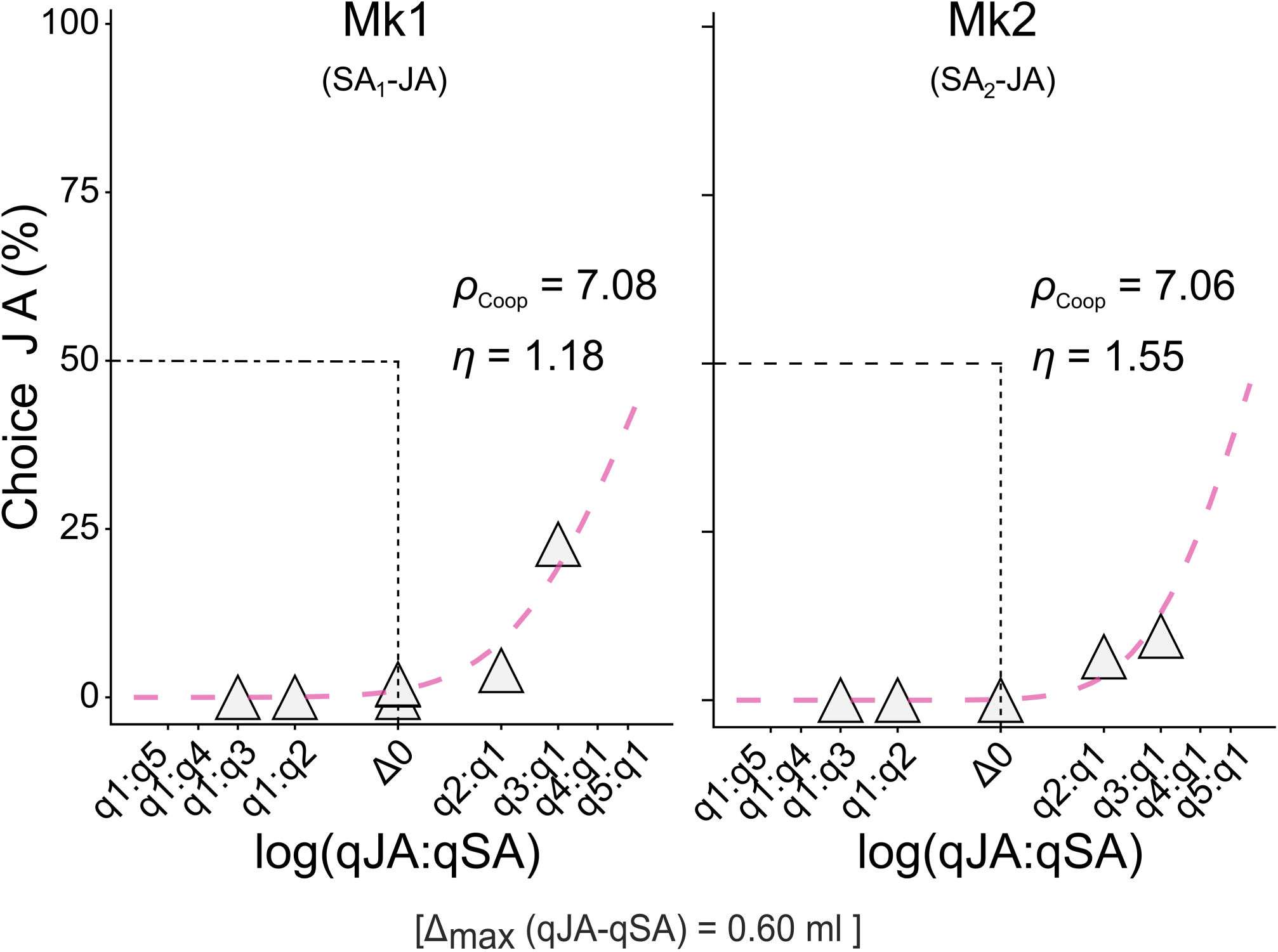
Cost-benefit evaluation when monkeys decide to cooperate or not (*prePhase* sessions). Animal behavior was examined in a preliminary phase (*prePhase*) and an experimental phase (*expPhase*) using tasks with identical trial structures, but differing in the reward magnitudes associated with the two offers (see Methods). Plots refer to aggregated data from the *prePhase* sessions, in which the maximal reward difference between each pair of offers was 0.60 ml. The percentage of choices in favor of cooperation is plotted on the y-axis against the log quantity ratio between the two offers for JA vs SA (x-axis). Each data point represents percentages of ‘JA chosen’ trials for one offer pair, across sessions. were obtained from probit regressions. The relative value (ρ_Coop_) and sigmoid steepness (η) are reported as text. Note how the choices are starkly biased in favor of the SA regardless of the relative difference between offers.

**Figure S4.**
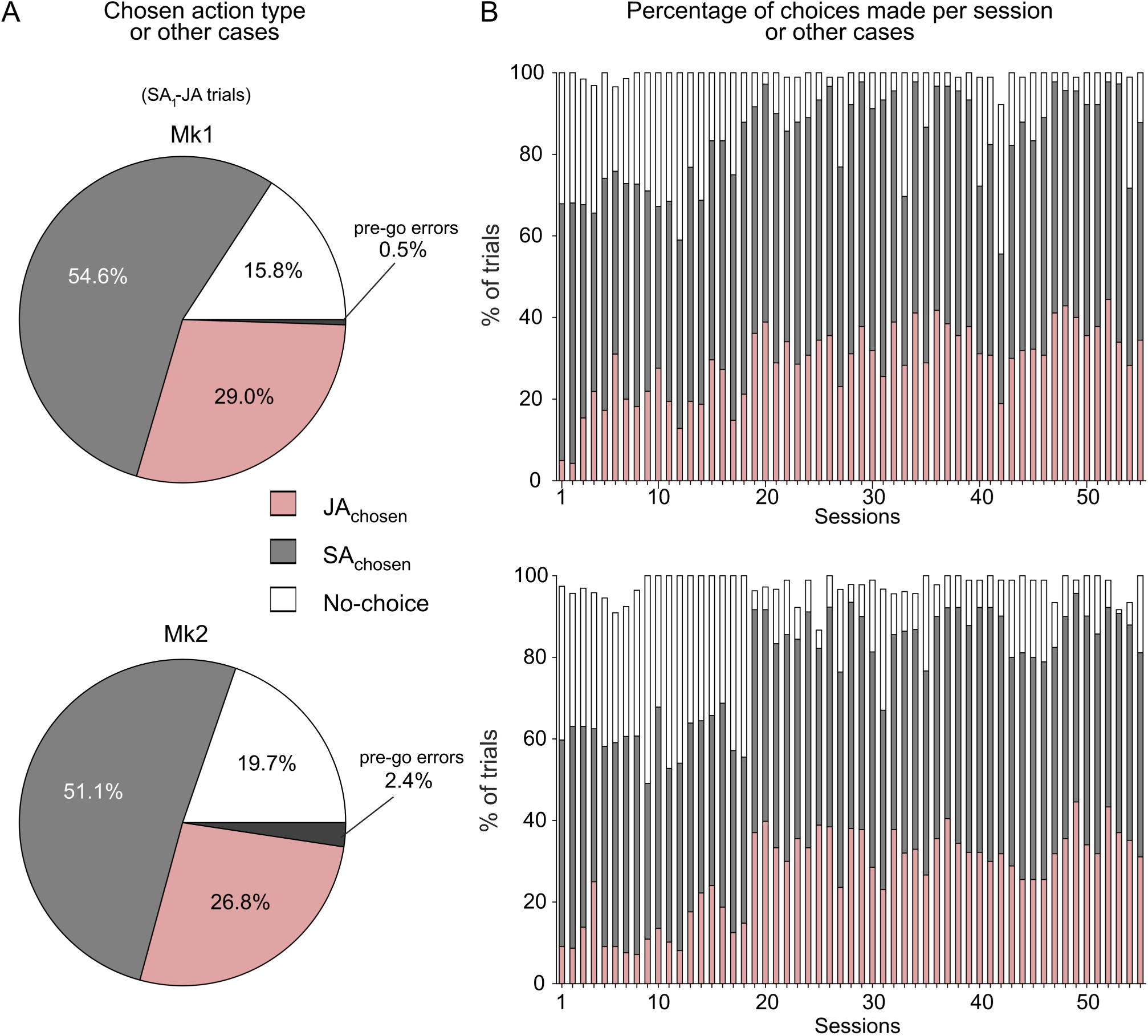
Distribution of choice responses in SAᵢ–JA trials. **(A)** Pie charts show, for SAᵢ–JAᵢ trials, the proportion of each choice outcome for the two monkeys when acting as ‘decider*’*: cooperative (JA, pink), individual (SA, gray), or decision avoidance (No-choice, white). A small fraction of trials (0.5% for Mk1 and 2.4% for Mk2) resulted in premature responses before the go signal (pre-GO errors). **(B)** Distribution of choices across sessions, showing the progressive decrease of decision avoidance trials over time, corresponding to an increase of choices in favor of JA. Similar distributions were observed for both monkeys.

